# Phages-bacteria interactions underlying the dynamics of polyhydroxyalkanoates-producing mixed microbial cultures via meta-omics study

**DOI:** 10.1101/2024.12.25.630301

**Authors:** Jian Yao, Yan Zeng, Xia Hong, Meng Wang, Quan Zhang, Yating Chen, Min Gou, Zi-Yuan Xia, Yue-Qin Tang

**Author notes:** Correspondence: Yue-Qin Tang.

## Abstract

The dynamics of the structure of polyhydroxyalkanoates-producing mixed microbial cultures (PHA-MMCs) during enrichment and maintenance is an unsolved problem. The effect of phages has been proposed as a cause of dynamic changes in community structure, but evidence is lacking. To address this question, we enriched five PHA-MMCs and sampled temporally to study the interactions between phage and prokaryotic members by combining metagenomics and metatranscriptomics. 963 metagenome-assemble genomes and 4294 phage operational taxonomic units were assembled from bulk metagenomic data. There were complex interactions between the phages in Casadabanvirus and Unclassified Hendrixvirinae and the dominant species in *Azomonas*, *Paracoccus*, *Thauera* and *Breundimonas*. The dynamic change of the structure of phage and prokaryotic communities were remarkably consistent. Further structural equation modeling analysis showed that phage communities could significantly affect the activity and community structure of prokaryotic microorganisms. In addition, seven key auxiliary metabolic genes (phaC, fadJ, acs, ackA, phbB, acdAB and fadD) in the PHA synthesis pathway were identified from phage sequences. Importantly, these auxiliary metabolic genes were expressed at the transcriptional level, indicating that they were in an active functional state. This meta-analysis provides the first catalog of phages in PHA-MMCs and the auxiliary metabolic genes they carry, as well as how they play a role in the dynamic changes of prokaryotic communities. This study provides a reference for subsequent studies on understanding and regulating the microbial community structure of artificial open microbial systems.

**Importance:** The synthesis of biodegradable plastic PHA from organic waste through mixed microbial cultures (MMCs), with extremely low cost, has the potential for expanded production. However, there is a gap of understanding of the dynamics of dominant species in MMCs to better control community structure. Our results demonstrate for the first time the impact of phages on the structure of prokaryotic communities in the MMCs. There are complex interactions between the PHA producers (*Azomonas*, *Paracoccus*, *Thauera*) and phages (Casadabanvirus and Unclassified Hendrixvirinae). Phage communities can negatively regulate the activity and structure of prokaryotic communities. In addition, the auxiliary metabolic genes in the PHA synthesis pathways carried by phages may promote the PHA synthesis ability of prokaryotic members. This study highlights the impact of phages on prokaryotic community structure, suggesting that phages have the potential to become a tool for better controlling the microbial community structure of PHA-MMCs.

## Introduction

Mixed microbial cultures (MMCs), which use complex microbial communities composed of multiple microorganisms as biocatalysts, has achieved significant achievements in pollutant removal(1) and waste utilization(2). Polyhydroxyalkanoates (PHAs) production by MMCs is a prime example. Due to outstanding biodegradability and material properties, PHAs have the potential to replace petroleum-based plastics in the face of increasing plastic pollution and fossil resource depletion(3). The high cost of PHA production with pure culture techniques has limited the widespread use of PHAs materials(4). As a result, PHA production with MMCs techniques that can utilize inexpensive waste biomass, has been proposed and is more likely to facilitate commercial scale-up of PHA production due to its low production costs(4).

Currently, the process for the production of PHAs by MMCs using complex waste biomass has three main steps(5). The process includes: 1) acid production, producing short-chain carboxylic acids (SCCAs, such as acetate, propionate, butyrate, valerate and lactate) that served as substrates of PHAs production by anaerobic degradation; 2) MMCs enrichment, enriching microbial communities with PHAs-synthesizing capacity from activated sludge by applying selective pressure; 3) PHAs accumulation, the maximum PHAs accumulation of enriched MMCs using SCCAs. Of these, the second step is the most important in the process. Nowadays, aerobic dynamic feeding (ADF) is the main strategy to enrich MMCs(6). Alternating cycles of the feast (carbon rich) and famine (carbon poor) in the ADF provides selection pressure. Because, PHA producers can accumulate intracellular PHAs as a backup carbon source during the feast phase and are better able to gain a competitive advantage during the famine phase(7). However, the dominant species in enriched microbial communities continue to change even under stable maintenance conditions(7). Huang et al. hypothesize that this is due to the “kill the winner” effect of phages where virulent phages prey on dominant species, causing other species to gain a dominant position and resulting in repeated replacement of dominant species(7, 8). Unfortunately, there is still no evidence to support this hypothesis, and the effect of phages on microbial community assembly during the enrichment and maintenance of PHA-MMCs has not been studied.

In recent years, researchers have begun to uncover the effects of phages on the structure and function of microbial communities in natural environments (e.g., deep-sea sediment(9) and soil(10)) and engineered systems (e.g., municipal wastewater treatment plants(11-13)). Virulent phages hijack the hosts to produce progeny and eventually lyse the host, and host cell debris fuel other microorganisms, which can directly affect the structure of microbial communities(14, 15). In addition, both virulent and temperate phages can manipulate host metabolism by reprogramming host metabolic pathways or expressing auxiliary metabolic genes (AMGs) to influence the function of microbial communities. For example, in the ocean, many AMGs are involved in photosynthesis(16), the pentose phosphate pathway(17), nitrogen metabolism(18), and sulfur metabolism(19). Given that the enrichment of PHA-MMCs is aimed at increasing PHA synthesis ability in microorganisms, phages may play an important but yet undiscovered role by influencing the abundance of dominant species or encoding AMGs related to PHA synthesis. Therefore, further studies are needed to examine the phage-host relationship in PHA-MMCs and extra PHA metabolic capabilities conferred by AMGs in phages.

We constructed five PHA-MMCs enrichment reactors with different SCCAs as the sole carbon source and collected 30 biological samples over the time scale. Based on 30 PHA-MMCs metagenomes and 30 corresponding metatranscriptomes, we recovered the high-quality assembled phage contigs (APCs). Our study aimed to explore the following questions: (1) Identification of phages in PHA-MMCs and characterization of their interactions with bacteria; (2) Whether and how phages affect the structure of the bacterial community; (3) Whether phages encode PHA synthesis-related AMGs and whether these AMGs are transcriptionally active. We recovered 4293 APCs and used structural equation model (SEM) to discuss the effects of phages on the activity of the prokaryotic microorganisms and the microbial community structure. Host prediction of APC and prediction of host-encoded antiphage defense systems were performed to explore the characteristics of phage-host interactions during enrichment and maintenance processes. The AMGs involved in PHA synthesis were identified in phages and complementary metatranscriptome analysis was conducted to verify the active expression of AMGs, which supported new discovery of active phages-mediated PHA biosynthesis function.

It is the first study to elucidate the effect of phages on the community structure and function of PHA-MMCs. The results provide guidance for subsequent studies to construct efficient and robust PHA-MMCs.

## Results and discussion

### Overview of the prokaryotic and phage communities in the PHA-MMCs

To investigate phage diversity during the enrichment and maintenance process of PHA-MMCs, as well as the effect of phages on the microbial structure, we enriched five PHA-MMCs with acetate, propionate, butyrate, valerate, and lactate as the sole carbon source, respectively, and incubated the MMCs for a period of 145 days (Figure 1A). The PHA synthesizing capacity of the five PHA-MMCs was rapidly increased in the first 17 days (enrichment phase) and remained stable thereafter (maintenance phase). The butyrate-enriched PHA-MMCs had the highest PHA synthesizing capacity, followed by the acetate, lactate, and valerate-enriched PHA-MMCs, and the lowest being the propionate-enriched PHA-MMCs (Figure 2A).

**Figure 1.**
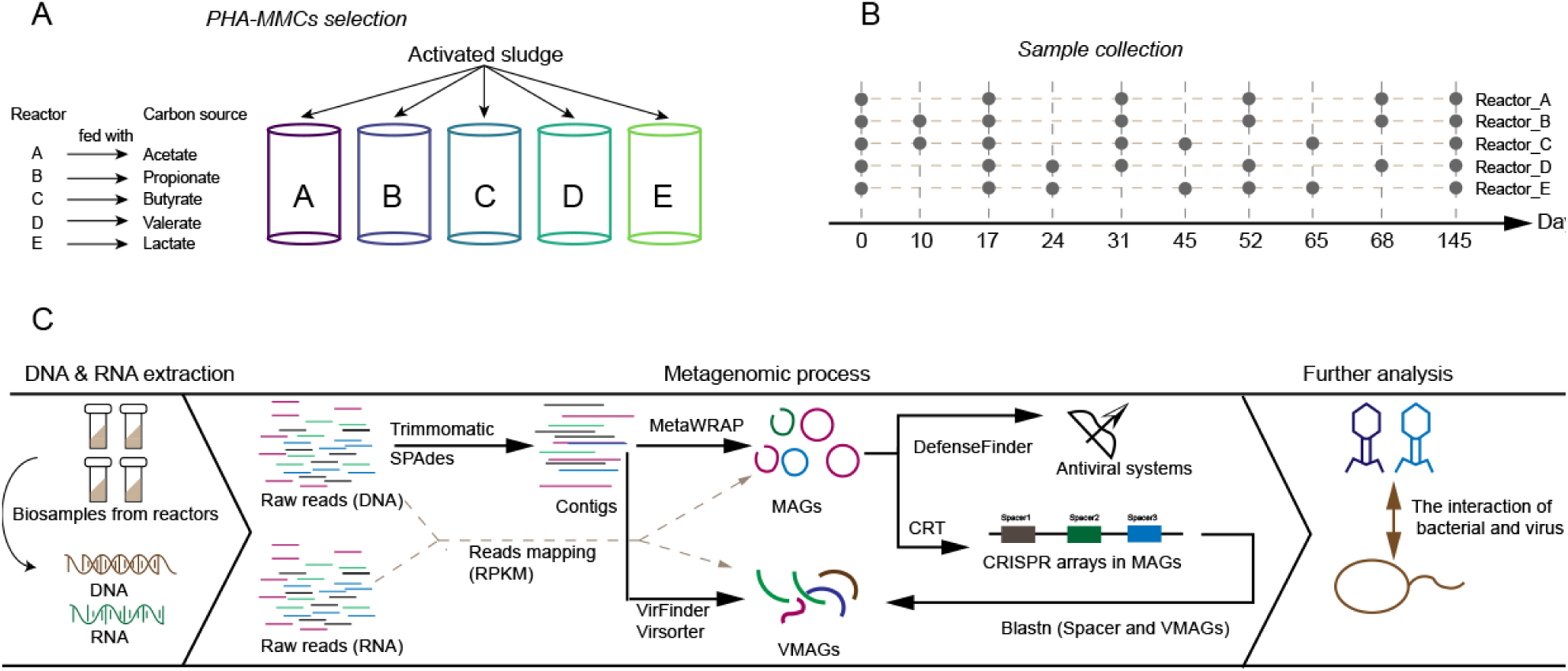
Experimental design and analytical processes. (A) The five enriched PHA-MMCs; (B) The distribution of biological samples collected; (C) The analytical processes of meta-omics data.

**Figure 2.**
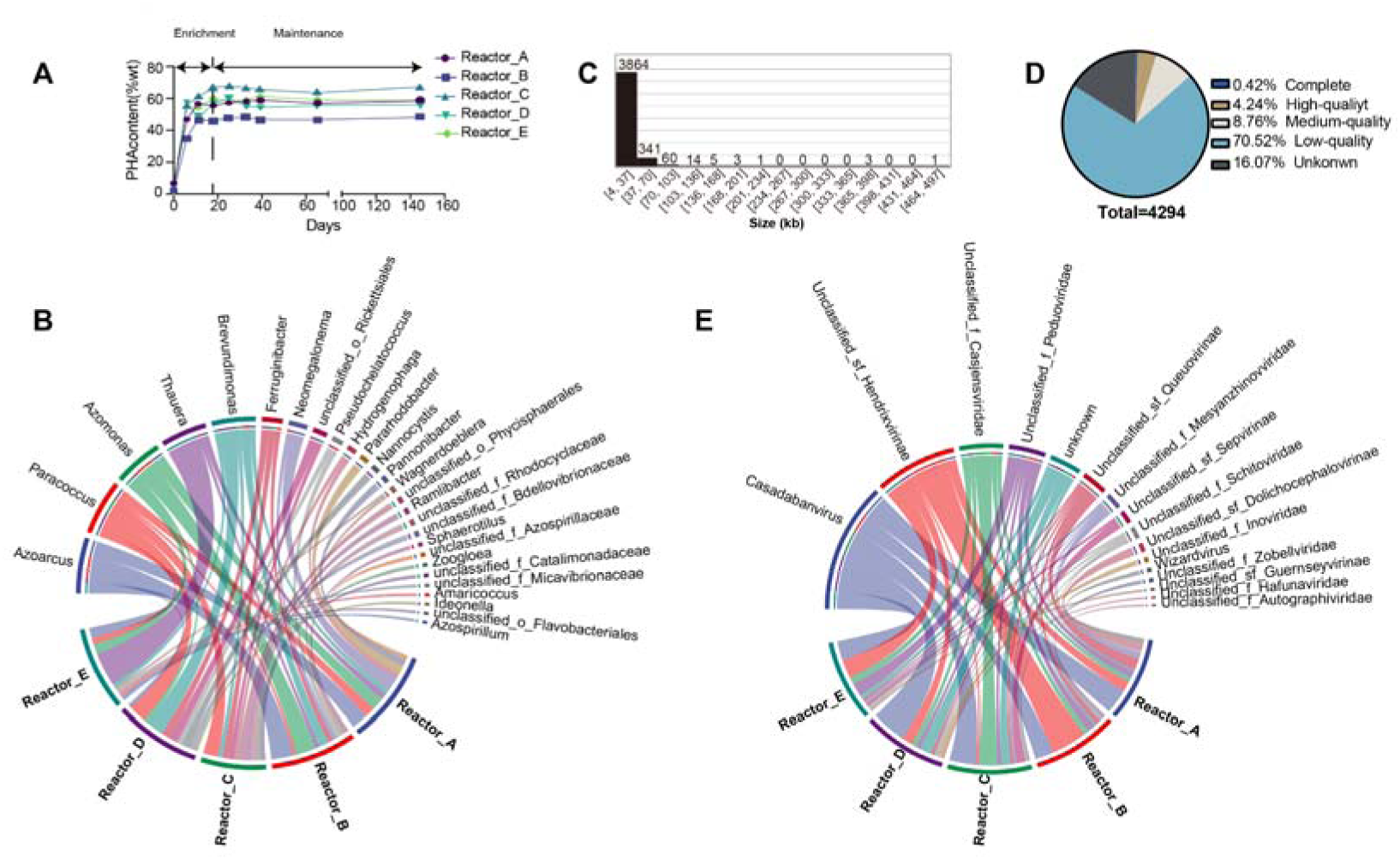
The overview of prokaryotic and phage communities. (A) The maximum PHA accumulation content (wt%) of PHA-MMCs in each reactor; (B) The top 10 bacterial genera (abundance) in each reactor; (C) The statistical histogram of pOTU size; (D) The percentage of pOTUs of different quality; (E) The top 10 pOTU genera in each reactor. The size of lines represents the abundance of the genus.

To further understand the dynamics of the prokaryotic and phage community structure of PHA-MMCs and the interactions between them, 30 time-sequenced samples were collected and subjected to meta-omics sequencing and analysis during the enrichment and maintenance of MMCs (Figure 1B, 1C). From these metagenomes, we recovered 963 high-quality metagenome-assemble genomes (MAGs) with completeness ≥ 70% and contamination ≤ 10% by de novo assembly and binning. After clustering by dRep(20), 711 unique MAGs were obtained, of which 112 were from Reactor_A, 144 from Reactor_B, 179 from Reactor_C, 141 from Reactor_D, and 135 from Reactor_E. Among five reactors, the bacterial MAGs belong to Proteobacteria (n=69, 86, 98, 93, and 83 in Reactor_A, B, C, D, and E, respectively) and Bacteroidota (n=25, 38, 46, 25, and 28 in Reactor_A, B, C, D, and E, respectively) dominated in the prokaryotic communities (Figure S2). The distribution of these MAGs in the different genera was shown in Figure S3-S7. In general, *Azoarcus*, *Paracoccus*, *Azomonas*, *Thauera,* and *Brevundimonas* were the dominant genus which was consistent with previous studies(7, 21, 22) (Figure 2B). However, the dominant genus was not the same in different reactors. For example, *Azoarcus* was dominant in Reactor_A and B, rather than in Reactor_C and D and *Thauera* dominated in Reactor_E, rather than in others, which might be due to the fact that different bacteria preferred different organic acids(21, 23-25).

The phage sequences were assembled and identified from bulk metagenomes containing sequences of diverse origins, which have been widely used to study viral communities in soil(26), deep-sea sediment(9), and deep-sea hydrothermal vents(27). In this study, VirSorter2(28) and VirFinder(29) were used to identify assemble phage contigs (APCs) from bulk metagenomes, resulting in 5719 putative APCs (Contigs ≥ 5 kb(30)). These APCs were then clustered at 95% identity and 80% coverage to generate 4294 phage operational taxonomic units (pOTUs). The size of these pOTUs ranged from 5001 to 496772 bp, in which six pOTUs were larger than 200 kb and possibly corresponded to giant phages (Figure 2C). From the quality assessment of pOTUs by checkv(31), we found 576 pOTUs (13.42%) were of medium quality and above, including 18 complete pOTUs (0.42%), high-quality pOTUs (4.24%) and medium-quality pOTUs (8.76%) (Figure 2D). VirFinder and VirSorter2 are powerful tools for identifying phage sequences from metagenomes and have been used in many studies(32). The low proportion of high-quality phage sequences also appeared in previous studies, which may be due to insufficient understanding of the extremely complex virosphere(11, 30). The VirFinder use the frequencies of DNA k-mers found in known viral genomes to train machine-learning classifiers to identify phage sequences(29). VirSorter2 use domain percentages, gene content features, and key homology genes in a tree-based machine learning framework to classify phage reads(28). Compared with VirFinder, VirSorter2 is more advanced in that it moves beyond a single model to represent the virosphere for phages identification. However, VirFinder and VirSorter2 are heavily dependent on existing virus databases. The incompleteness of existing bacteriophage databases limits the output of better-quality phages identification results.

The life type of pOTUs was predicted based on the protein composition and associations by PhaTYP(33). Among these pOTUs, 2276 pOTUs were classified as temperate phages (53%) while 1741 pOTUs were classified as virulent phages (40.5%) The average abundance of temperate phages identified in these biospecimens was 1.7 times that of virulent phages. This might be due to the fact it is conventionally enriched for genomes of temperate phages that integrate into host genomes in bulk metagenomic data(9). The lack of complete annotation of lytic-specific genes and the incomplete assembly of phage genomes also could lead to underestimation of potent phage signals.

The taxonomy of pOTUs was classified by PhaGCN2 which was a GCN-based model and learned the species masking feature via a deep learning classifier(34). Totally, 2007 pOTUs (46.7%) were classified at the phylum level, with the Uroviricota (n = 1902) as the most dominant phylum in each reactor (Figure S2). Zhang et al. studied the phages in activated sludge from three different sewage treatment plants, using three methods (PhaGCN2, CAT and BLASTn to the IMG/VR database) for taxonomic classification and the results also showed that Uroviricota was the most prevalent phylm(12). Some pOTUs can’t be classified, due in part to the fact that the current viral phylogeny is still understudied, which also occurs in other studies(30, 35). Among the Uroviricota, 51 genera were identified for pOTUs and the *Casadabanvirus* (n = 740, a genus in class Caudoviricetes) was the most dominant genera of the class Caudoviricetes, followed by the three unclassified genera in Hendrixvirinae (subfamily), Casjensviridae (family), and Peduoviridae (family) respectively (Figure 2E). This is consistent with the study of Ju et al. in which, phages in sludge from 32 wastewater treatment plants were taxonomically classified using vConTACT(11).

Consistent with the prokaryotic community, the abundance of members in the phage community changes dynamically over time (Table S1-S5). We used the mean rank shift (MRS)(36) to quantitatively characterize this dynamic change of phage and prokaryotic communities (Figure S8). MRS is a temporal analogue of species rank-abundance distributions and indicate the degree of species reordering between two time points. 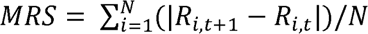. where *N* is the number of species in common in both time points, *t* is the time pointand *R_i_*_,*t*_ is the relative rank of species *i* at time *t*.

In these communities, the MRS of phages at adjacent time points ranged from 400 to 1000, which was much higher than that of bacterial members. This was partly because the number and diversity of phages far outstripped bacterial members in PHA-MMCs. In addition, there is a positive correlation between the MRS of phages and bacterial communities (Spearman’s Rho and p-value in Reactor A: 0.89 & 0.037; Reactor B: 0.65 & 0.1; Reactor C: 0.81 & 0.04; Reactor D: 0.87 & 0.02; Reactor E: 0.37 & 0.2). However, MRS did not consider the loss and gain of members during community succession. Therefore, more powerful evidences were needed to verify the impact of phages on prokaryotic community structure.

### Evidence that phages shape prokaryotic communities

To show the differences among the community structure of prokaryotic and phage communities in the different samples, the Bray-Curtis distance among samples was calculated and sorted by Principal Co-ordinates Analysis (PCoA) (Figure 3A, 3B). Sample points in each reactor did not cluster together during the late stages of maintenance (days 65, 68, and 145), which showed that the prokaryotic community structure remained in dynamic flux during maintenance, even when a deterministic selection pressure was applied (alternating satiation and starvation phases). In addition, the structure of the phage communities in different samples also did not converge.

**Figure 3.**
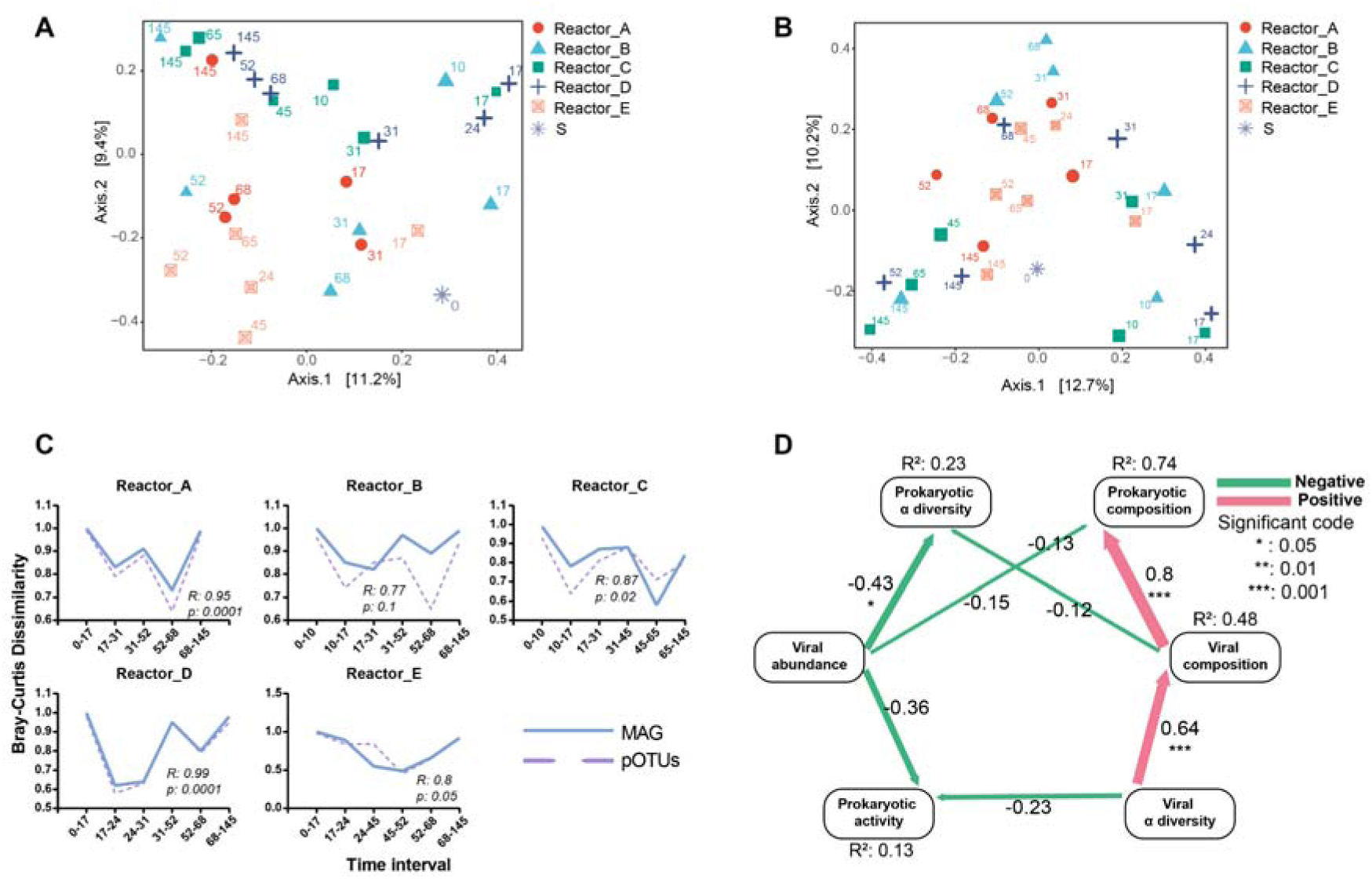
Temporal dynamics of prokaryotic and phage communities. (A, B): The PCoA of prokaryotic communities (A) and phage communities (B). The “S” denotes activated sludge inoculum and the numbers indicate the days of sampling; (C): The Bray-Curtis community dissimilarity though days for prokaryotic (solid line) and phage (dashed line) communities. The dissimilarity illustrates the change in community composition from the previous time point. Pearson’s correlation test was tested between the dissimilarity of prokaryotic and viral communities; (D): Path diagram of SEM showing the effect of phage communities on prokaryotic communities. Composition is represented by the PC1 from the Bray-Curtis dissimilarity-based principal coordinate analyses. The α diversity is represented by the Chao1 index. Numbers adjacent to the arrows are standardized path coefficients (r). “*” means the significant index (Chi-squared test) p≤0.05 and “***” means the significant index p≤0.001. R^2^ represents the proportion of variance explained for every dependent variable in the model.

To investigate whether the dynamics of prokaryotic community structure could be mirrored in phage communities, we quantified these temporal changes in community structure by Bray-Curtis values and calculated the Spearman’s Rho between prokaryotic and phage communities (Figure 3C). The Bray-Curtis values described the degree of difference between the community structures of temporally adjacent samples; the larger the Bray-Curtis value, the greater the difference. There was a significant positive relationship between the prokaryotic and phage communities shifts and their dynamics mirrored well (Spearman’s Rho in Reactor_A, B, C, D & E = 0.95, 0.77, 0.87, 0.99 & 0.8). This tended to be caused by the interaction between phage and bacteria. Under the selective pressure of enrichment, PHA producers gradually gained a competitive advantage, leading to the increased abundance in the community. The encounter rate with the corresponding phage would be greatly increased by an increase in host abundance, which promoted the proliferation of phages(37, 38). The virulent phage hijacked hosts for self-replication and then lysed the host cells. Under phage predation, the number of hosts gradually decreased, which negatively affected phage proliferation, leading to a subsequent decrease in the abundance of phages. However, more evidence is needed to prove phage predation on hosts, which is the key to showing that phage is one of the driving forces in changing the structures of prokaryotic communities described above.

To solve this problem, we used structural equation modeling (SEM) to discern the causality and quantify the effects of the drivers (Figure 3D). The significant positive correlation (r: 0.8, p ≤ 0.001) between phage community composition and prokaryotic community composition was consistent with the above analyses (Figure 3C), as phages were dependent on the host for survival. Importantly, there was a significant negative correlation (r: -0.43, p ≤0.05) between phage abundance and the alpha diversity (Chao1) index of the prokaryotic community, in addition to a negative correlation (r: -0.36) with prokaryotic community activity. This might indicate that virulent phages hijacked and killed host cells during proliferation, thereby reducing the activity of the prokaryotic hosts. As the phage proliferated and lysed host cells, the number of hosts rapidly decreased, which in turn reduced the Chao1 index of the prokaryotic community. This further suggested that the synergistic changes between the phage and prokaryotic communities were not caused by the response of the temperate phages to the dynamics of hosts, as they benefit from the high growth rate of the host while limiting their lytic activity(37, 39). In summary, we added important evidence for the hypothesis that phages shape the structure of prokaryotic community in PHA-MMCs through the “kill the winners” effect.

This result might inspire us to develop a method to use phage to in situ regulate the microbial community structure of PHA-MMCs to improve their PHA synthesis efficiency. The efficiency of PHA synthesis by PHA-MMCs was lower than that of pure bacterial fermentation, partly because there were members in its microbial community that did not synthesize PHA, resulting in substrate waste(40). Unfortunately, the selection pressure applied by the current bacterial enrichment strategy could not eliminate these non-PHA producers. If we could isolate and culture phages that can specifically lyse these non-PHA producers, and add these phages into PHA-MMCs to specifically eliminate non-PHA producers. In theory, we could obtain PHA-MMCs with higher PHA synthesis efficiency.

### Phage-host interaction dynamics

To specifically elucidate the interactions between phages and hosts, we screened the putative phage-hosts linkages and the antiviral defense systems of hosts. During the interaction between phages and bacteria, bacteria use the CRISPR-Cas system to add spacers to their own genome to resist the next infection(41). This allows us to search for potential host-phage relationships by comparing spacer sequences in bacteria with pOTU sequences. Overall, we recovered a total of 16725 spacers from all 30 metagenomes. All five reactors had a positive relationship between the number of spacers recovered and time (Figure S9 A). In addition, we observed that the number of spacers also had a significant and strong positive relationship with the total number of unique pOTUs across all samples (Figure S9 B). These results suggested that the CRISPR-Cas system played an important role in the ongoing interaction between bacteria and phages and integrated matching spacers into CRISPR arrays through time.

Furthermore, we searched for hosts of 200 high-quality pOTUs using CRISPR-match and genomic homology matching. 1421 linkages were detected between 129 pOTUs (including 15 genera) and 409 hosts (including 73 genera) (Figure 4A). We constructed the linkages between phages and bacterial hosts as interaction networks at the genus level. This interaction network had 88 nodes and 198 links. The statistical characteristics of each node in the interaction network can be found in Table S6. Among these nodes, 61 nodes have a degree greater than 1. This result might indicate that the phage-host linkage was not unique, but rather that a single phage had the potential to infect multiple hosts, or that a single host was infested by multiple phages, which was consistent with previous studies in oil wells(42), earthworm intestines(30), and acid mines(43). For example, the hosts of the phage genus *Casadabanvirus* were distributed among 47 bacterial genera, including PHA-synthesizing bacteria such as *Azoarcus*, *Hydrogenophaga*, and *Paracoccus*. Secondly, there tended to be a significant positive correlation between the abundance (of both phage and host) and the number of phage-host linkages at the genus level (Figure S10). One reason for this might be that a high abundance of hosts that adapted to selective pressures (e.g., PHA producers) provided high encounter rates for potential phages, and sufficient hosts also made these phages much more abundant than others(37). Another possible reason was that linkages between high-abundance phages and the host were more easily detected, as certain rare phages might not even be recovered and assembled with high quality. This dynamic linkages between phages and bacteria might have a potential role in maintaining community stability. These phages might selectively infect and lyse community members that were overabundant in the community, such as (*Azoarcus* and *Paracoccus*), providing niches for other community members and preventing a single member from dominating the microbial community(37). This emergent state in which community members took turns to assume the leading position in the community had higher biodiversity and community resilience, leading to better community stability(44).

**Figure 4.**
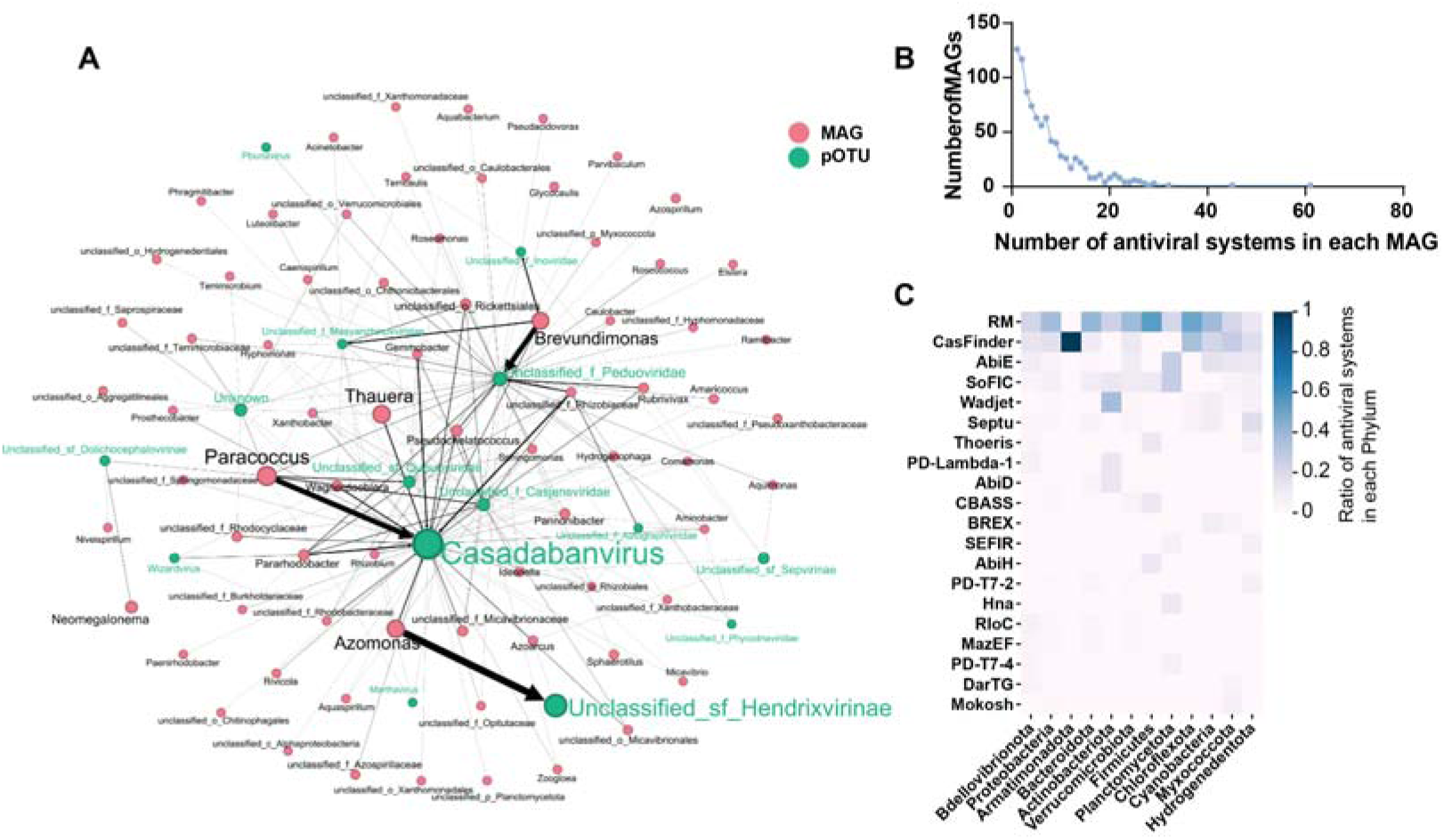
The interaction between phages and hosts. (A) The interaction network of phages and hosts. The red nodes and green nodes represent the hosts and phages. The size of nodes represents the abundance of genus. The size of edges represents the number of linkages between phages and hosts. The “unclassified_f_Casjensviridae” represents the unclassified genus in the family Casjensviridae. The “Unclassified_sf_Sepvirinae” represents the unclassified genus in the subfamily “Sepvirinae”. (B) The histogram of the number of antiviral systems in each host. (C) The proportion of top 20 antiviral systems in each host phylum.

In the co-evolution of phage and host, the hosts also adapted different antiviral systems to resist phage infection(45). We used DefenseFinder to discover the antiviral systems encoded in the prokaryotic MAGs and found that the antiviral arsenal of the prokaryotes was highly variable which was consistent with Tesson’s study(45). Overall, 86% of the MAGs encoded two or more types of antiviral weapons (Figure 4B). However, as the number of encoded antiviral systems increased, the number of corresponding MAGs decreased rapidly, suggesting that encoding more antiviral systems was not a perfect survival strategy. Of the 109 antiviral systems detected, RM and CRISPR-Cas were dominant, accounting for 36.5% and 14.6% of the total, respectively (Figure 4C). The RM and CRISPR-Cas provide defense through the degradation of viral nucleic acids, which appear to be the most widespread antiviral systems used by prokaryotes in a variety of habitats(30, 45, 46). The RM and CRISPR-Cas systems were the predominant antiviral systems in 10 of the 12 bacterial phyla, while Wadjet and SoFIC were the predominant antiviral systems in Actinobacteriota and Planctomycetota, respectively. In addition, hosts defended themselves against phages through various abortive infection systems, which were ‘altruistic’ cell death systems that were activated by phage infection and limited viral replication(46). For example, some members of Cyanobacteria, Myxococcota, and Hydrogenedentota had the AbiE antiviral system which were encoded by bicistronic operons and functioned via a non-interacting (Type IV) bacteriostatic TA mechanism(47). Members of Firmicutes had Thoeris antiviral system. When triggered by phage infection, the ThsB sensor of the Thoeris defence systems produced signalling molecules, which activated the NAD^+^ degrading activity of ThsA, leading to the depletion of the host NAD^+^ pool and abortive infection(48). This may indicate that there were differences in the antiviral strategies of different prokaryotes.

To specifically characterize the temporal dynamics of the phage-host linkages in each reactor, we used network diagrams to show the phages associated with the host at different time periods (Figure S11-S15). We calculated the sequence similarity among the corresponding phages of each host using VIRIDIC(49). The results showed that most of the phages had very low sequence similarity to each other’s nucleotide sequence, suggesting that the prokaryotic members of PHA-MMCs could be infected by diverse phages (Table S7). The linkages of the host and phages changed dynamically at different time points. For example, in Reactor_D (Figure S14), the linkages of *Paracoccus* between phage 0111B24 were detected on days 52 and 145, but not on day 68. The loss of linkages or the appearance of new linkages also happened on the *Paracoccus* in Reactor_E (Figure S15) and the *Thauera* in Reactor_A (Figure S11). However, there were also many “strong phages” that form tight linkages with the host and were detected at all times, such as the phage 1010E_19 with *Brevundimonas* in Reactor_A (Figure S11). Among these hosts, *Azomonas* (in Reactor_B, C & E) and *Brevundimonas* (in Reactor_A & D) had a more solid linkage with the phage, even though they were in different reactors. While, the phage infecting *Paracoccus* (in Reactor_C, D & E) and *Thaurea* (in Reactor_A) were more variable at different time points. Furthermore, phage characteristics (e.g. lifestyle) were not significantly related to the diversity of these linkages. This indicated that the stability of host-phage linkages was more dependent on host characteristics.

### Potential impacts of AMGs on the metabolisms and the PHA synthesis of host

Phages can influence biogeochemical processes in ocean(9), municipal wastewater treatment plants(11), and gut(30) by delivering auxiliary metabolic genes while infecting hosts. In a recent study, researchers found that phages in paddy soil carried a large number of auxiliary metabolic genes related to carbon fixation. In addition, in situ isotopic labeling experiments induced by mitomycin-C revealed that ^13^CO_2_ emissions from the treatment with added lysogenic phage decreased by approximately 17.9%(50). In addition, Ju et al. analyzed the phages in the sludge of 23 sewage treatment plants and found that these phages carried a large number of AMGs involved in carbon (acpP and prsA), nitrogen (amoC), sulfur (cysH) and phosphorus (phoH) metabolism, and these AMGs were active at the transcriptional level(11).

To investigate the effect of phages on the PHA anabolism of PHA-MMCs, AMGs encoded in pOTUs were predicted using DRAMv pipelines and functionally annotated by the KEGG database. These AMGs were generally involved in four types of activities, namely metabolism, environmental information processing, genetic informational processing and cellular processes (Figure S16). The high proportion of AMGs related to purine metabolism, pyrimidine metabolism, amino acid biosynthesis, mismatch repair, homologous recombination, and DNA replication might be due to their favorability for phage replication in the host, which was consistent with other studies(9, 11). It was worth noting that some AMGs in fatty acid metabolism may be involved in the biosynthetic pathway of SCCAs (acetate, propionate, butyrate, valerate, and lactate) into PHA, which needs further identification.

Overall, these SCCAs were converted into (R)-3-hydroxybutyryl-CoA and (R)-3-hydroxyvaleryl-CoA through a series of biochemical reactions, and then polymerized into poly(3-hydroxybutyrate) (PHB) and poly(3-hydroxyvalerate) (PHV) by PHA synthase. After the conversion of acetate and lactate to acetyl-CoA, two molecules of acetyl-CoA were condensed to acetoacetyl-CoA by beta-ketothiolase. The acetoacetyl-CoA was reduced to (R)-3-hydroxybutyryl-CoA by acetoacetyl-CoA reductase. Butyrate and valerate were converted to (R)-3-hydroxybutyryl-CoA and (R)-3-hydroxyvaleryl-CoA by beta-oxidation pathway, respectively. As for propionate, it was firstly converted to propionyl-CoA. Together with acetyl-CoA, they served as substrates for beta-ketothiolase, followed by the reduction of 3-ketovaleryl-CoA to (R)-3-hydroxyvaleryl-CoA.

Seven key A MGs involved in PHA anabolism were further screened and identified from the functionally annotated AMGs using the NCBI CD-search tool. These AMGs were distributed among 10 phages and 24 hosts (Figure 5A). The *phaC* was the most important AMG, as it encodes a PHA polymerase that polymerizes the precursors (R)-3-hydroxybutanoyl-CoA and (R)-3-hydroxyvaleryl-CoA into poly(3-hydroxybutyrate) (PHB) and poly(3-hydroxyvalerate) (PHV)(51). The genes *fadJ* (3-hydroxybutanoyl-CoA epimerase)(52) and *phbB* (acetoacetyl-CoA reductase)(53) catalyze the isomerization of (S)-3-hydroxybutanotyl-CoA and the reduction of acetoacetyl-CoA, respectively, to produce (R)-3-hydroxybutanoyl-CoA, providing precursors for PHA synthesis. In addition, *acs* (acetyl-CoA synthetase)/ackA (acetate kinase)(54, 55), *acdAB* (acetyl-CoA synthetase)(56), and *fadD* (long-chain fatty acid-CoA ligase)(57) catalyze the conversion of acetate, propionate, and valerate to acetyl-CoA, propionyl-CoA, and valeryl-CoA, respectively, facilitating the entry of organic acids into the PHA synthesis pathway (Figure 5B).

**Figure 5.**
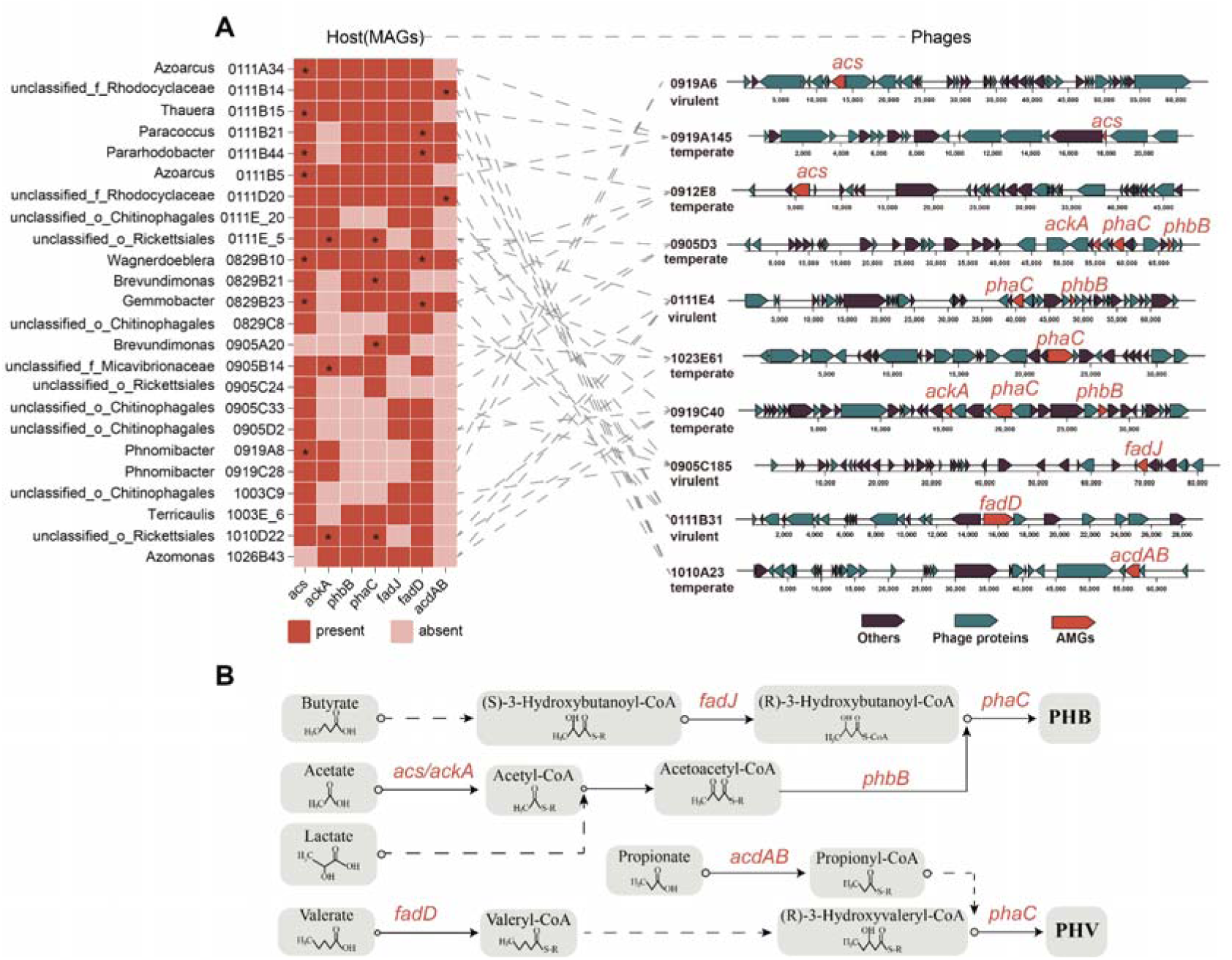
The AMGs involved in PHA anabolism. (A) The distribution of AMGs in phages and hosts. The dotted line indicates the linkages between hosts and phages. The “**\***” indicates a high similarity (≥80%) between the host and phage AMGs. “unclassified_f_X” represents the unclassified genus in family X. “unclassified_o_X” represents the unclassified genus in order X. (B) Metabolic function of AMGs in PHA synthesis. The dotted lines represent the multi-metabolic pathways.

Furthermore, we found that the AMGs in phage and host were diverse in sequence by phylogenetic analysis and sequence similarity calculation (Figure S17-S23). For example, the gene *phaC* annotated as PHA synthase in hosts of a phage was not the same as AMG in this phage (Figure S17). The *phaC* of phage 0905D3 was very similar to the nucleotide sequence in host YJ0111E.5, but significantly different from that in another host YJ0905C.24. Additionally, *phaC* of 0905D3 had transcriptional activity in sample 0905C (Table S8). These phenomena were also present for the genes of *fadD*, *acdAB*, *acs*, and *ackA* (Figure S18, S20, S22, S23). Unexpectedly, in the case of both *phbB* and *fadJ*, the sequence similarity between the AMGs in the phage and host was less than 75% (Figure S19, 21). However, the fact that these AMGs carried by phages were transcriptionally active in different samples still implied that phages were able to transfer AMGs among hosts and enhance PHA anabolism in the hosts.

## Conclusions

In this study, we investigated the dynamic interaction between phages and bacterial members in PHA-MMCs. In the PHA-MMCs system, phages could infect and lyse dominant species to vacate ecological niches for other species, resulting in a community succession state in which dominant species alternated. This might retain a high species diversity and maintain community stability. This provides a potential explanation for understanding the emergence of community members in open artificial microbial systems. In addition, while infecting different hosts, phages might express auxiliary metabolic genes related to PHA anabolism to enhance the PHA anabolism capacity of community members. Of course, the above conclusions need to be verified by further phage supplementation experiments. Therefore, in subsequent studies, the in-situ regulation of phages on PHA-MMCs can be considered to enhance its effectiveness in practical engineering applications. One is “phage therapy”, which can be used to add relevant phages to eliminate members in PHA-MMCs that do not have the ability to synthesize PHA to enhance the utilization efficiency of carbon sources. The second is to add phages carrying key genes in the PHA synthesis pathway to enhance the PHA synthesis capacity of community members.

## Materials and Methods

### PHA-MMCs enrichment

Five sequencing batch reactors (SBR), each with a working volume of 2 L, were used for the cultivation of PHA-MMCs. Activated sludge from a municipal wastewater treatment plant (Chengdu, China) was used as the inoculum source. Activated sludge was collected from the aerobic stage of the Anaerobic-Anoxic-Oxic process and filtered to remove solid impurities (screen size of 0.25mm). Reactors A, B, C, D, and E were fed with synthetic wastewater containing acetate, propionate, butyrate, valerate, and lactate as the sole carbon source, respectively (Figure 1A). The content of inorganic components and trace elements in synthetic wastewater is described in supplementary information S1. Five SBRs were operated according to the ADF selection strategy, with a cycle length of 12 h, consisting of feeding (10 min), aeration (650 min), settling (50 min), and discharge (10 min). All reactors were operated at 26°C. More detailed information can be found in supplementary information S1 and Figure S1.

The maximum PHA accumulation content (wt%) of PHA-MMCs at different times was evaluated by fed-batch PHA accumulation assays. The dissolved oxygen (DO) concentration was maintained at above 4 mg/L by aeration. The DO meter (WTW, Germany) was used to detect the DO concentration in the fermentation broth. The 200ml fermentation liquid in enrichment reactor was transferred to a new 400ml reaction vessel, and then 200ml of fresh substrate was added. Air was provided to the reactor through an aeration pump. After aeration for 1 hour, aeration was stopped. After standing for 15 minutes, 200ml of supernatant was removed. Subsequently, 200ml fresh substrate was added to the 400ml vessel and aerated for the second batch fermentation. After 1 hour of reaction, the above steps of stopping aeration, standing and removing supernatant were repeated. The above operation was repeated for another three times. After the fifth fermentation, 10ml of fermentation liquid was taken and centrifuged at 10000rpm and10min to collect the biomass. The biomass was pre-treated and the PHA content of biomass was determined using GC-MS. The detailed determination method was described in supplementary information S2.

### Sample collection, DNA/RNA extraction and sequencing

During the 145 days of the SBR operation, ten time points were selected for biological sampling (Figure 1B). The fermentation broth was collected in sterile DNase- and RNase-free centrifuge tubes and centrifuged at 12,000 rpm for 30 min at 4°C. The biomass was collected and stored in liquid nitrogen. Total DNA and RNA were extracted via the cetyl-trimethyl ammonium bromide (CTAB) method(58). DNA and RNA sequencing were performed on Illumina NovaSeq (Illumina Inc., San Diego, CA, USA) at Majorbio Bio-Pharm Technology Co., Ltd. (Shanghai, China). The size of each metagenomic and metatranscriptomic raw sequence data was more than 15 GB and 9 GB, respectively. Sequence data associated have been deposited in the NCBI Sequence Read Archive database (PRJNA1121485, the accession numbers could be found in table S9). Details of library construction and sequencing were described in supplementary information S3. The Trimmomatic v0.36(59) was used to process raw DNA and RNA sequencing data to get clean raw data (ILLUMINACLIP: adapters/TruSeq3-PE.fa:2:30:10; LEADING:3; TRAILING:3; SLIDINGWINDOW:6:30; MINLEN:100).

### Bacterial metagenome-assembled genomes (MAGs) assembly and bioinformatics analyses

The clean DNA reads from 30 metagenomes were spliced into contigs by SPAdes v.3.5.0(60). The bacterial MAGs were assembled from contigs by MetaWRAP v.1.2.1(61). For subsequent analysis, high-quality MAGs (completeness ≥ 70% and contamination ≤ 10%) were selected. The taxonomy of MAGs was classified by GTDB-Tk version 2.1.1 based on the Genome Taxonomy Database(62). The metagenomic and metatranscriptomic reads were mapped to MAGs by BBMap (v35.85; http://sourceforge.net/projects/bbmap/). The abundance and activity of MAGs were calculated as reads per kilobase transcript per million reads (RPKM).

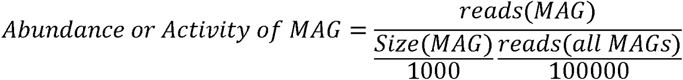

In the equation, the “reads (MAG)” is the number of reads of metagenomic (or metatranscriptomic) raw data mapped to each MAG. The “reads (all MAGs)” is the sum of reads of all MAGs. The “size (MAG)” is the number of base pairs of each MAG. To compare differences in microbial communities at different enrichment times, all MAGs from the same reactor were clustered using dRep v.2.6.2 (-pa 0.9, -sa 0.99)(20). The abundance and activity values of clustered MAGs at different time points were summarized as abundance or activity (Table S1-S5) for subsequent statistical analysis. The CRISPR spacers were searched from MAGs scaffolds by the CRISPR recognition tool (CRT, v2.1)(63). The antiphage systems of MAGs were detected by DefenseFinder v.1.2.2(45).

### Assemble phage contig (APCs) identification and bioinformatics analyses

The APCs were identified comprehensively from metagenomic contigs by VirSorter2 v.2.1.0(28) and VirFinder v1.1(29). A contig (length ≥ 5 kb) that met one of the following three conditions was selected as an APC, (i) VirSorter v2.1 score ≥0.9; (ii) VirFinder v1.1 score ≥ 0.9 along with p < 0.05; (iii) both VirSorter2 score ≥0.5 and VirFinder ≥ 0.7, p < 0.05. CD-HIT v.4.7 was used to cluster the APCs to obtain phage operational taxonomic units (pOTUs) with parameter -c at 0.95 and -aS at 0.8. The abundance of pOTUs was calculated as reads per kilobase transcript per million reads (RPKM) by BBMap (v35.85; http://sourceforge.net/projects/bbmap/).

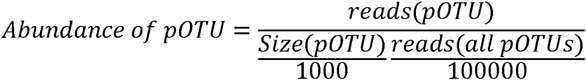

In the equation, the “reads (pOTU)” is the number of reads of metagenomic raw data mapped to each pOTU. The “reads (all pOTUs)” is the sum of reads of all pOTUs. The “size(pOTU)” is the number of base pairs of each pOTU. The abundance of pOTUs in five SBRs were described in Table S1-S5. The CheckV v.1.0.3(31) was used to assess the quality of pOTUs (completeness ≥ 90%; contamination ≤ 60%). The taxonomy of pOTUs was classified by PhaGCN2 v.2.0(34). The lifestyle of pOTUs was predicted by PhaTYP(33) function in PhaBOX(64).

The auxiliary metabolic genes (AMGs) of pOTUs were identified and annotated by DRAM-V v.1.5.0(65) pipeline as follows: The VirSorter2 v.2.1.0(28) (--prep-for-dramv) was run on pOTUs sequences, and then the AMGs were predicted from the resulting sequences by DRAM-V v.1.5.0(65). The putative AMGs were further confirmed using the NCBI CD-search tool (https://www.ncbi.nlm.nih.gov/Structure/cdd/wrpsb.cgi) with a threshold value of e-value < 10^−5^.

The phage-host linkages prediction was performed on two different in silico strategies: (i) CRISPR-spacers match. The CRISPR spacers of MAGs were queried for exact matches to pOTUs sequences by BLASTn v.2.9.0(66). Only the matches (identity ≥ 97%, coverage ≥ 90%, mismatch ≤ 1) were regarded as highly confident phage-host linkages. (ii) The sequence homology between MAGs and pOTUs. The sequences of pOTUs were compared with MAGs by BLASTn v.2.9.0(66). The matches (identity ≥ 70%, bit score ≥ 50, alignment length ≥2500, e-value ≤ 0.001) were regarded as highly confident phage-host linkages.

### Statistical analysis and visualization

The α diversity indexes and principal coordinates analysis (PCoA) of phage and microbial community were calculated in R v.4.3.2 with package “phyloseq”(67). Based on the abundance table of community members, the R package “vegan” was used to calculate the Bray-Curtis dissimilarity between bacterial and phage communities at adjacent time points to characterize the degree of change in community member structure(42). The Spearman correlation coefficient and p-value between the Bray-Curtis dissimilarity of bacterial and phage communities, were provided by the function cor.test() in the R v.4.3.2. To further analyze the impact of phage communities on bacterial communities, structural equation modeling (SEM) was constructed with phage community characteristics (α diversity, abundance, and activity) as predictor variables and bacterial community characteristics(α diversity, abundance, and activity) as response variables. The SEM analysis was performed R v.4.3.2 with the package “piecewiseSEM” v.2.3.0 (https://jslefche.github.io/piecewiseSEM/). The network diagram was visualized by Gephi v.0.10. The heatmap, phylogenetic tree, and gene cluster map were visualized by chiplot (https://www.chiplot.online/).

## Supporting information

supplement_informations

## Acknowledgment

This study was funded by the China Petroleum and Chemical Corporation (No. 421063-10).

## Data availability

Sequence data associated have been deposited in the NCBI Sequence Read Archive database (PRJNA1121485).

## Author contributions

Y.T. and J.Y. conceived the project anddesigned the experiments. J.Y. did the experi-ments, analyzed data, and wrote the manuscript. Y.T. supervised the project and revised the manuscript. Y.Z., Y.C., M.G. and Z.X. contributed to the methodology. X.H., M.W. and Q.Z. contributed to sample collection and nucleic acid extraction.

## Ethics statement

No animals or humans were involved in this study.

